# Tbx5 overexpression in embryoid bodies increases TAK1 expression but does not enhance the differentiation of sinoatrial node cardiomyocytes

**DOI:** 10.1101/2023.03.04.531127

**Authors:** Yunkai Dai, Fatemeh Nasehi, Charles D. Winchester, Ann C. Foley

**Affiliations:** Clemson University, Department of Bioengineering, 68 President Street, Charleston, SC

**Keywords:** Sinoatrial node, SAN, Tbx5, TAK1, Embryoid Body

## Abstract

Genetic studies place Tbx5 at the apex of the sinoatrial node (SAN) transcriptional program. To understand its role in SAN differentiation, clonal embryonic stem (ES) cell lines were made that conditionally overexpress Tbx5, Tbx3, Tbx18, Shox2, Islet-1 and Map3k7/TAK1. and cardiac cells differentiated using embryoid bodies (EBs). EBs overexpressing Tbx5, Islet1, and TAK1 beat faster than cardiac cells differentiated from control ES cell lines suggesting possible roles in SAN differentiation. Tbx5 overexpressing EBs showed increased expression of TAK1, but cardiomyocytes did not differentiate as SAN cells. They showed no increase in the expression of Shox2, or Islet1 and decreased expression of HCN4. EBs constitutively overexpressing TAK1 direct cardiac differentiation to the SAN fate, but also have decreased phosphorylation of its targets, p38, and Jnk. This opens the possibility that blocking the phosphorylation of TAK1 targets may have the same impact as forced overexpression. To test this, we treated EBs with 5z-7-Oxozeanol (OXO), an inhibitor of TAK1 phosphorylation. Like TAK1 overexpressing cardiac cells, cardiomyocytes differentiated in the presence of OXO beat faster and showed increased expression of SAN genes (Shox2, HCN4, and Islet1). This suggests that activation of the SAN transcriptional network can be accomplished by blocking the phosphorylation of TAK1.

## Introduction

The SAN, which initiates and propagates electrical impulses to coordinate contractions in the heart is an anatomically discrete region of specialized myocardium. The transcription factor Tbx5, has been a primary target for studies of SAN differentiation since it was first linked to Holt-Oram syndrome, a disease characterized by cardiac conduction system abnormalities (Basson et al. 1994, Holt and Oram 1960, Newbury-Ecob et al. 1996).

Tbx5 is a T-box of transcription factor (Basson et al. 1997, Li et al. 1997) which plays a critical role in the specification and maintenance of the conduction system by regulating the conduction system transcriptome. (Arnolds et al. 2012, Moskowitz I. P. et al. 2007, Moskowitz Ivan PG et al. 2004). Germline deletion of Tbx5 in mice leads to embryonic lethality by E10.5 and severely impaired SAN differentiation (Bruneau et al. 2001). The atrioventricular conduction system fails to mature in Tbx5 haploinsufficient (Tbx5^-/+^) mice resulting in prolongation of the PQ interval (Moskowitz Ivan PG et al. 2004). Together these data indicate that Tbx5 is necessary for the differentiation of the conduction system generally and SAN differentiation specifically.

A transcriptional program for SAN differentiation has been defined through genetic studies, placing Tbx5 at the apex(Christoffels et al. 2010). Overexpression of transcription factors downstream of Tbx5 in pluripotent stem cells has been shown to activate the differentiation of cardiomyocytes with some, or all, of the characteristics of SAN cells (Ionta et al. 2015, Jung J. J. et al. 2014, Kapoor et al. 2013) however the exact role of Tbx5 in these studies is unclear. In addition, these data and ours presented below suggest that induction of specific cardiac fates by transcription factor overexpression may be highly dose-dependent. The differentiation of SAN-like cells from pluripotent cells can also be activated by manipulating intracellular signaling cascades mediated by BMP (Protze et al. 2017) and Map kinase signaling (Brown et al. 2017, Wiese et al. 2011). It is not clear how manipulation of signaling pathways impacts the SAN transcriptional program. We previously showed that constitutive overexpression of the Map kinase Map3k7/TAK1 impacts this program but notable impacts the expression of Tbx5 and Nk2.5 very early during EB differentiation with Nkx2.5 expression being repressed and Tbx5 being upregulated earlier than any other cardiac transcription factor that we assessed (Brown et al. 2017). To test the importance of Tbx5 we developed constructs to overexpress Tbx5 and other members of the SAN transcriptional program under the control of doxycycline (DOX) inducible promoters. Tbx5 overexpression did cause cardiomyocytes derived from EBs to beat faster but did not activate other signs of SAN differentiation however these EBs did surprisingly overexpress TAK1. Constitutive overexpression of TAK1 directed cardiomyocyte differentiation to the SAN fate but also repressed the phosphorylation of TAK1 downstream targets including p38 and Jnk (Hunter et al. 2019). Since the constitutive overexpression of TAK1 represses the phosphorylation of known downstream targets, it is possible that blocking TAK1 phosphorylation might have the same effect as forced overexpression. To test this EBs were treated with 5z-7-oxozeanol (OXO), an inhibitor of TAK1 phosphorylation. Cardiac cells differentiated in the presence of OXO beat faster, showed morphological changes, expressed the SAN channel protein HCN4 and expressed key SAN transcription factors Shox2 and Islet1, supporting this hypothesis.

## Materials and Methods

### Cell culture and Embryoid Body (EB) Differentiation

R1 mouse embryonic stem cells (ATCC) were transduced with aMHC::GFP to track cardiac differentiation (Brown et al. 2010). These cells were subsequently transduced with lentiviruses encoding DOX-inducible-overexpression cassettes for Tbx5 (pTripZ-mTbx5), Tbx3 (pTripZ-mTbx3), Tbx18(pTripZ-mTbx18, Shox2 (pTripZ-mShox2), Islet1 (pTripZ-mIsl1), and Map3k7/TAK1 (pTripZ-mTAK1). Clonal lines were isolated and individually verified for expression of genes of interest. Map3k7/TAK1-overexpressing mESCs was previously described (Brown et al. 2017). These mESCs were maintained in standard growth medium and kept under constant selection with puromyocin to maintain cells with continued expression of the pTripz-lentiviral constructs (2 mg/ml). For EB differentiation, these mESCs were passaged off of feeder cells (Mouse embryonic fibroblasts) and differentiated as EBs using the hanging drop method, as previously described (Brown et al. 2010). At appropriate days, EBs were treated with either DOX diluted in DMSO or DMSO alone.

### Construction of the pTripZ-mTbx5, DOX-inducible-Tbx5-overexpression vector

The open reading frames of mouse Tbx5, Tbx3, Tbx18, Shox2 or TAk1 were amplified by PCR and directionally cloned downstream of the TurboRFP reporter using ClaI and MluI restriction sites in the pTripZ vector (Addgene) viral constructs were then validated by sequencing. Lentiviruses were produced using the second-generation lentiviral expression system (Bajpai and Terskikh 2007).

### Real-time quantitative PCR (qRT-PCR)

Approximately 100,000 ES cells were collected for each condition or time point analyzed. Total RNA was extracted with RNeasy Mini Kit, and 200 ng was used for first strand cDNA synthesis using QuantiTect Reverse Transcription Kit. qRT-PCR was performed with SybrGreen Master Mix, using 40 ng template/reaction on a Roche LightCycler® 480 Real-Time PCR Instrument, and analyzed with the LightCycler 480 software package. Crossing point data was first adjusted to reflect the efficiency of primer pairs by comparison to standard curves (based on dilution series over a total dynamic range of 1:1,000 or 1:10,000 for positive control cDNAs) and subsequently normalized to the ubiquitously expressed transcript GAPDH. Data represents averages ± standard error of 3 independent experiments. Further analysis was carried out using GraphPad Prism. Primers used in this study are as follows:

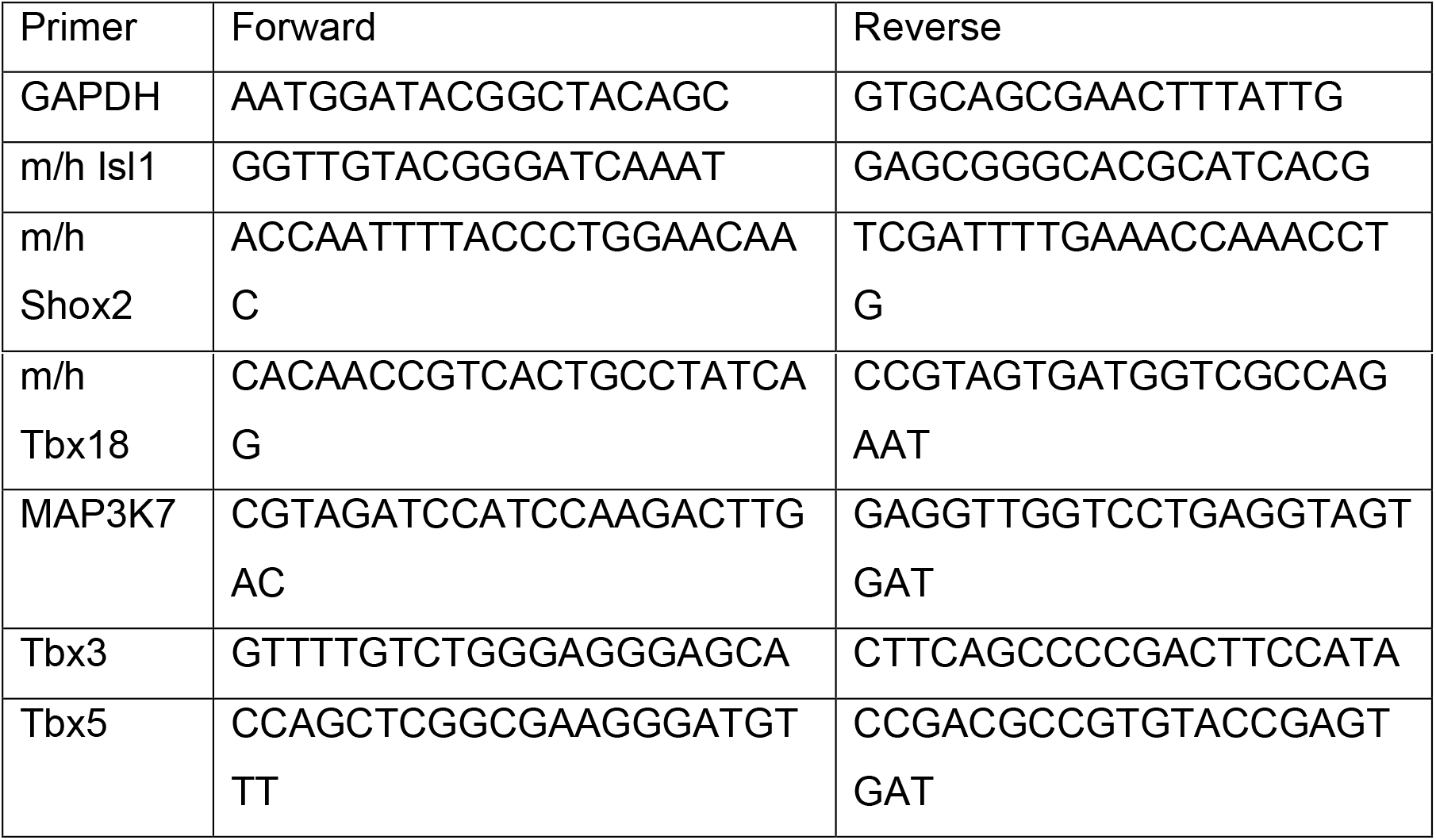

### Flow Cytometry

EBs were dissociated with 0.25% Trypsin/EDTA at 37 °C for 30 minutes filter through a 100 μm sieve to remove remaining undissociated cells and debris, and resuspended in FACS buffer (PBS plus 1% BSA and 10 ng/ml DNAse). Cells were counted using a hemocytometer, checking single-cell suspension at the same time. Flow Cytometry was performed with The Beckman Coulter MoFlo Astrios EQ cell sorter and data was analyzed using FlowJo VX software. Cardiac cells were sorted based on the expression of GFP driven by the aMHC promoter, as previously described (Brown et al. 2010).

### Immunocytochemistry

EB dissociation was performed as described above for Flow Cytometry. Cells were replated onto gelatin-coated 4-well chamber slides and allowed to attach for 18-24 hours. Slides were fixed in 4% paraformaldehyde for 15 minutes, gently washed 3 times for 5 minutes in PBS, blocked with either cytosolic antibody staining buffer (CASB) (PBS, 1% FBS, 0.1% BSA, and 0.1% TritonX-100) or nuclear antibody staining buffer (NASB) (1% FBS, 0.2% BSA and 0.25% Triton-X-100) for an hour. Cells were incubated with primary antibodies diluted in blocking buffer overnight. Cells were washed with PBS 3 times for 5 minutes and then incubated with Alexa Fluor-labeled secondary antibodies for an hour at room temperature. Three additional PBS washes were performed with DAPI added to the second wash. Coverslips were added and sealed with acrylic. Unless otherwise stated in the text, images were obtained using a Zeiss AxioImager microscopy. The primary antibodies were used as follow: Cx43 (Sigma, cat#C6219, 1:1000), CT3 (DSHB, cat#CT3, 1:250), DsRed (Clontech, cat#632392, 1:500), GFP (Thermo, cat#A11122, 1:500), HCN4 (Sigma, cat#SAB520035, 1:1000), Shox2 (Abcam, cat#ab55740, 1:1000), Tbx5 (Thermo, cat#42-6500, 1:500), Islet1 (Abcam, cat#ab20670. 1:500).

### Bradford Assay and Western Blots

Protein was extracted from different cell lines in RIPA buffer with protease inhibitors and sodium orthovanadate. Total protein was quantified by Bradford Assay (Pierce Coomassie Protein Assay Kit) compared to a serial dilution of BSA in a range of 0.064 - 2mg/ml. For western blots, 20 *μ*g/protein was loaded per well into prepared SDS-PAGE gels (Invitrogen NuPAGE 12% Bis-Tris gels). Proteins were transferred to PVDF membranes and immunoblotted for visualization with Immobilon Western chemiluminescent HRP substrate (Millipore) using a GeneGnome XRQ-chemiluminescence imaging system. Quantification was based on raw peak volume data. Antibodies used were anti-Tbx5 (Thermo, cat#42-6500, 1:500), TAK1 (Sigma, cat# AB1305414, 1:1000), HCN4 (Invitrogen, cat# MA3-903, 1:1000), Cx43 (Sigma, cat#C6219, 1:1000). Statistical significance was determined by student t-test with p<0.05 considered significant.

### Beat Rate

During EB differentiation, live cell imaging was used to determine the rate of beating for cardiomyocytes (identified based on the expression of αMHC::GFP). Beat rate was determined by manually counting beats per minute.

## Results

Six mouse ES cell lines that conditionally overexpress Tbx5, Islet1, TAK1, Shox2, Tbx18 and Tbx3 were generated using a lentivirus that drives gene expression and a fluorescent reporter under the control of a DOX-inducible promoter. This vector also has a selection cassette with puromycin resistance driven by the ubiquitin promoter (TRE::turboRFP-GOI;UBC::rtTA3-IRES-PURO) (Fig.1A). Expression of target genes was confirmed in clonal ES cell lines after 72 hours of DOX treatment by qRT-PCR (Fig.1B).

**Figure 1:**
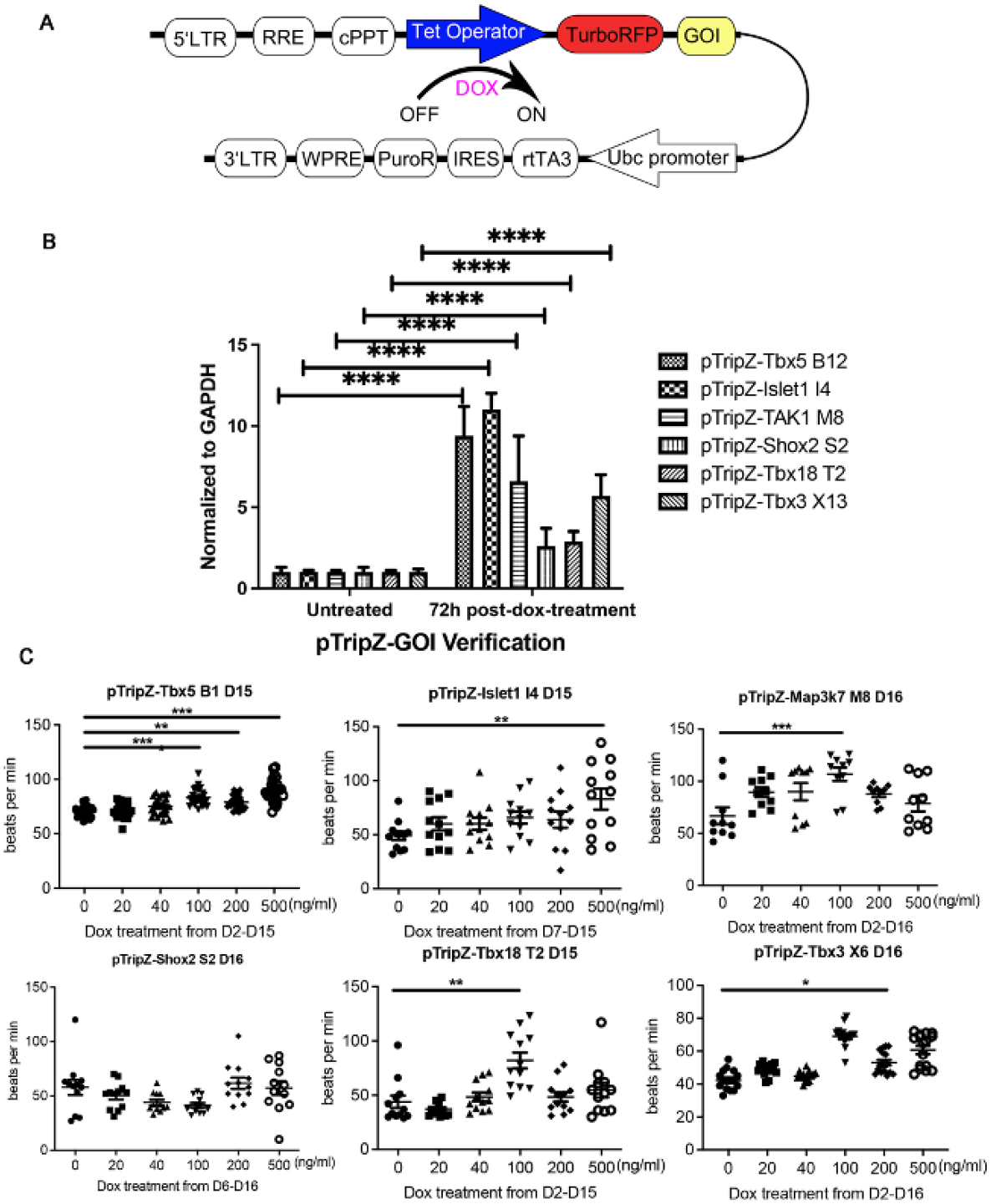
Clonal Isolation and testing of ES cell lines that conditionally overexpress important SAN genes. A) Schematic illustration of the inducible overexpression vecto r. B) QRT-PCR data of Tbx5, Islet1, TAK1, Shox2, Tbx18, and Tbx3, 72 hours post-Dox treatment compared to untreated cells. Data represent means ± standard error of 3 independent experiments. Statistical significance was determined by unpai red, two-tailed t-test. C) Manual beat data counting of Tbx5, Islet1, TAK1, Shox2, Tbx18 and Tbx3 overexpressing EBs at day 15 with DOX doses of 0ng/ml, 20ng/ml, 40ng/ml, 100ng/ml, 200ng/ml and 500ng/ml. Statisitical significance was determine by ANOVA. *p< 0.05, **p<0.01, ***p<0.001, ****p<0.0001.

As an initial test of each cell line’s potential to differentiate into SAN cardiomyocytes, mES cells were differentiated into EBs and treated with increasing doses of DOX (0-500ng/mL). In most cases treatment began at day 2 however, in two cases, Shox2 and Islet1, DOX treatment was started on day 6 and 7, respectively, based on their expression in wild type ES cells(Brown et al. 2017). Earlier treatment of these lines with DOX inhibited cardiac differentiation. The resulting EBs were evaluated for beat rate, an early indicator of SAN differentiation on day 15 or 16. Beating areas were identified based on visual inspection and confirmed by expression of aMHC::GFP which we previously showed to be a faithful reporter of cardiac differentiation (Brown et al., 2010).

Tbx5 overexpressing EBs had increased beat rate at 100 ng/mL, 200 ng/mL, and 500 ng/mL, while Islet1 overexpressing EBs showed a modest increase at 100 ng/mL and 200 ng/mL. Tbx3, Tbx18, and Map3k7 overexpressing EBs showed a significant increase at 100 ng/mL. Shox2 overexpressing EBs showed no change in beat rate across different doses.

Four independent clonal mES cell lines (B1, B5, B10 and B12) overexpressing Tbx5 were established, all showing stable, inducible upregulation of Tbx5 transcripts in ES cells, as confirmed by qRT-PCR at 24, 48 and 72 hours after addition of DOX (Fig.2A). Cardiomyocytes differentiated for 20 days from three of the four cell lines showed increased beat rate at one or more dosage of DOX (Fig.2B). The fourth clonal line showed only poor lineage differentiation and was excluded from further analysis. Of the remaining clones B1 cells showed the most robust cardiac differentiation and were therefore used for all subsequent analyses. Upregulation of TBX5 protein in response to DOX was confirmed in the B1 line by immunocytochemistry (Fig.1C).

**Figure 2:**
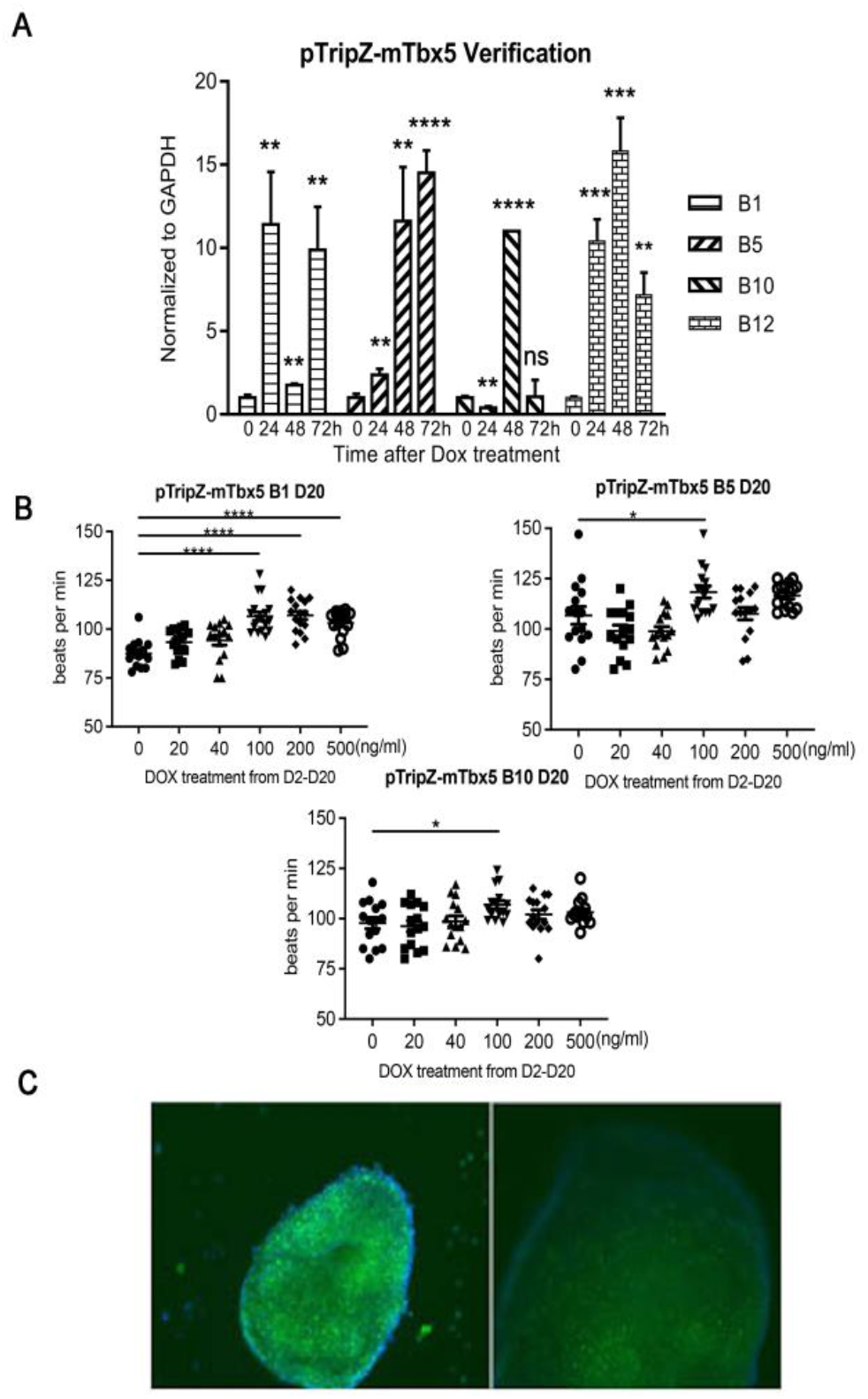
Verification of pTripZ-mTbx5; αMHC::GFP mouse ES cell lines. A) B1, B5, B10, B12 mES cells were treated with 1 μg /mL DOX and qRT-PCR was performed to detect relative expression of Tbx5 at 0, 24, 48, 72h. Statistical significance was determined by unpaired two-tailed t-test. B) Manual beat data counting of B1, B5 and B10 EBs at day 20 with DOX doses of 0ng/ml, 20ng/ml, 40ng/ml, 100ng/ml, 200ng/ml and 500ng/ml. Statistical significance was determined by ANOVA. C) Immunocytochemistry using anti-Tbx5 antibody in B1 mES cells 48h after DOX or DMSO addition. *p< 0.05, **p<0.01, ***p<0.001, ****p<0.0001.

### B1 ES cells show increased cardiac differentiation not related to the overexpression of Tbx5

Our previous experience studying cardiogenesis using ES cells suggests that clonal isolation can impact the cardiogenic potential of cells. To test this, EBs were differentiated with or without the addition of DOX from day 4 to day 21 (Fig.3). Cells were assessed on days 15, 17, 19, and 21 by flow cytometry for cardiac differentiation. Cardiomyocytes were identified based on the expression of GFP from the aMHC::GFP transgene and cardiac differentiation expressed as the percentage of GFP-expressing cells. The B1 line, with or without the addition of DOX showed increased cardiac differentiation as compared to the parent cell line (R1). These data suggest that the increase in cardiac differentiation in B1 EBs is due to clonal differences but was not impacted by Tbx5 overexpression.

**Figure 3:**
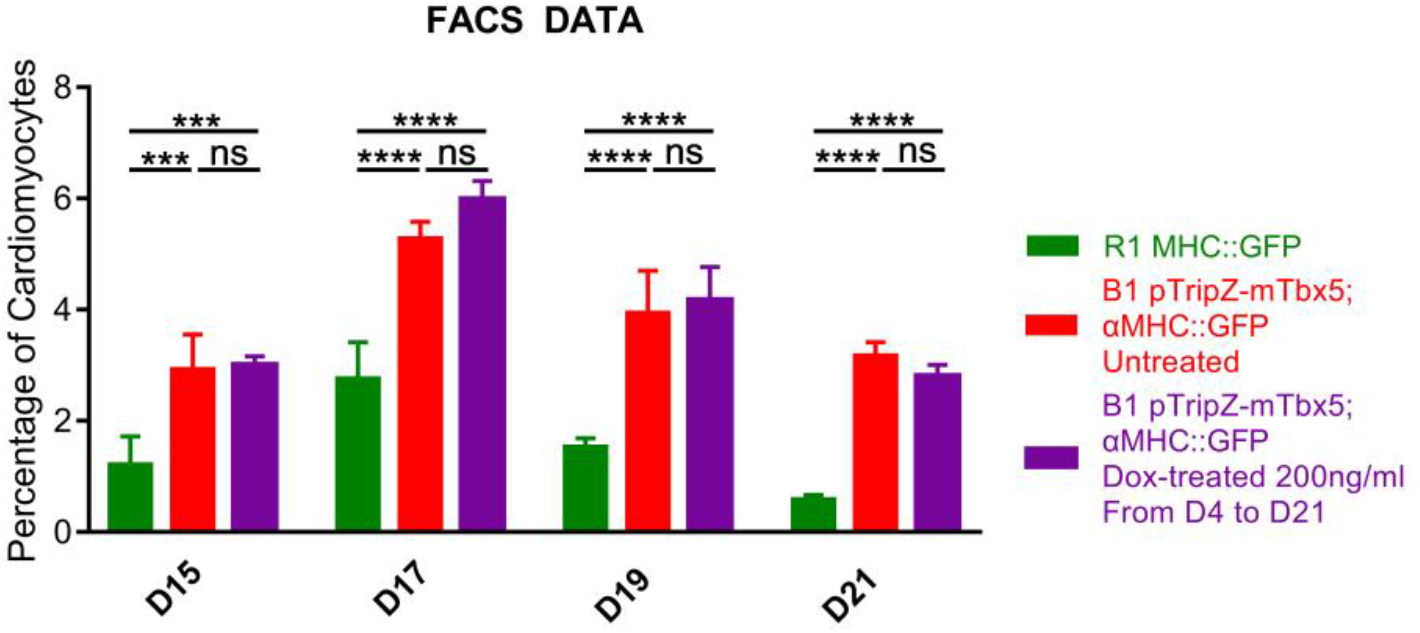
Cardiac differentiation analyzed by Flow Cytometry. The percentage of cardiomyocytes at day 21 of EB differentiation based on the number of cells expressing the aMHC::GFP reporter as a percentage of total cells by flow cytometry. Data represent means ± standard error of 3 independent experiments. Statistical significance was determined by unpaired, two-tailed t-test. ***p<0.001, ****p<0.0001.

### Day 21 Analysis of protein expression

At 21 days, DOX-treated EBs continued to overexpress Tbx5, as assessed by western blot (Fig.4). They also overexpressed TAK1. Interestingly, they showed downregulated overexpress the SAN marker HCN4. Protein expression of Cx43, which is expressed in all cardiac cells except SAN, was unchanged.

**Figure 4:**
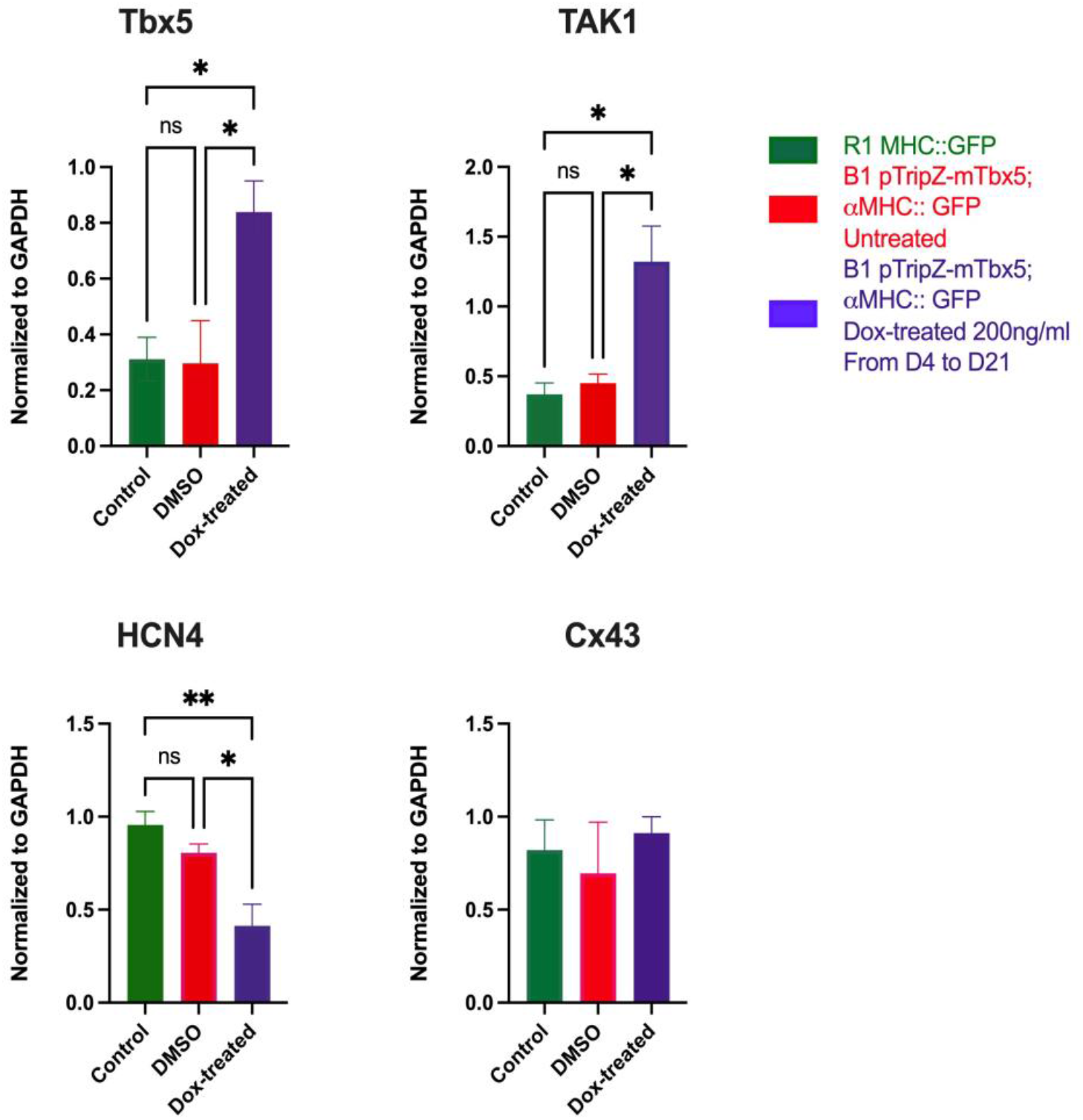
Relative protein expressions of R1, B1, B1 DOX-treated EBs at day 21. Tbx5, Map3k7/TAK1, HCN4 and Cx43 proteins were assessed by western blot. Data represent means ± standard error of 3 independent experiments, normalized to GAPDH. Statistical significance was determined by unpaired two-tailed t-test. *p<0.05, **p<0.01.

### Subtype of Differentiated Cardiomyocytes

Since cardiac cell represent only a fraction of the cells that differentiated in these EBs, we assessed the expression of SAN markers specifically in cardiomyocytes by immunocytochemistry. EBs were dissociated and individual cells replated in slide wells. Cardiac cells were identified using either expression of cardiac promoter reporters (aMHC::GFP or αMHC:mCherry) or by immunocytochemistry using the anti-cardiac troponin antibody (anti-CT3). Individual cardiomyocytes were immunostained for expression of HCN4, Shox2 or Connexin 43 (Cx43) (Fig.5A).

**Figure 5:**
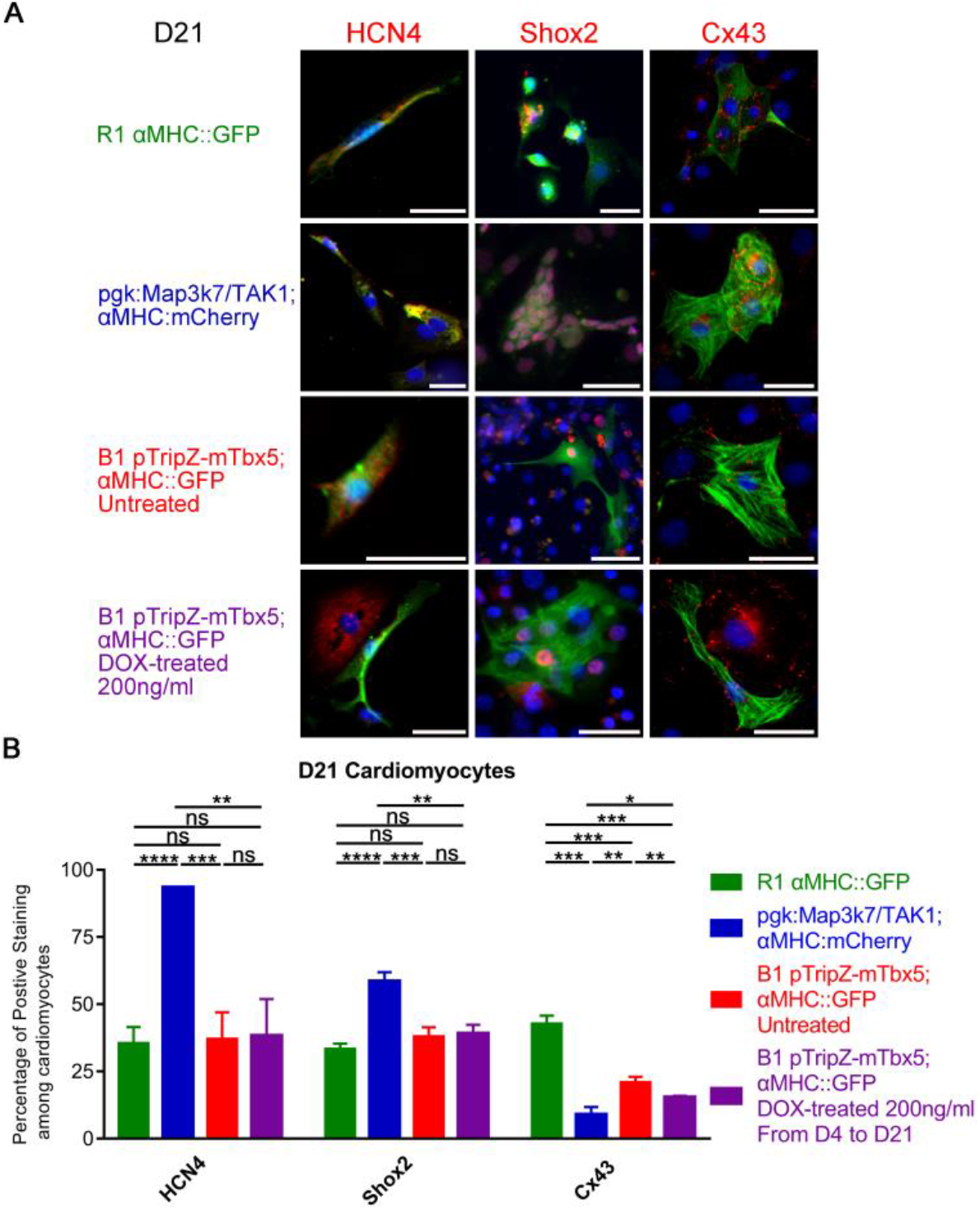
SAN marker expression in D21 cardiomyocytes. A) Representative image of ICC staining of day 21 cardiomyocytes of R1, Map3k7/TAK1-overexpressing, B1, and B1 DOX-treated EBs. EBs were dissociated into single-cell suspension and replated onto chamber slides. Then these cells were stained with GFP, DsRed or CT3 antibody to identified cardiomyocytes. HCN4, Shox2 and Cx43 antibody were immunostained with red fluorophore, DAPI labelled nuclei blue. Scale bar, 30 μm. B) The percentage of HCN4, Shox2 and Cx43 positive cells. Data represent means ± standard error of at least 3 independent experiments. Statistical significance was determined by unpaired two-tailed t-test. *p<0.05, **p<0.01, ***p<0.001, ****p<0.0001.

For these studies, we used B1 cells with or without the addition of DOX and positive and negative control lines including the parent cell line (R1; αMHC:GFP), and the Map3k7/TAK1-overexpressing cell line (PGK:Map3k7/TAK1; αMHC:mCherry) which directs cardiac cell differentiation to the SAN lineage (Brown et al. 2017). As expected, a higher percentage of cardiac cells derived from the TAK1 EBs expressed HCN4 and Shox2, whereas a lower percentage of these cells expressed Cx43, as compared to the R1 parent cell line (Fig.5B).

Neither the B1 nor the B1 DOX-treated EBs showed increased expression of HCN4 or Shox2, although these cells did show decreased expression of Cx43, similar to the TAK1 EBs (Fig.5B) but this did not depend on the addition of DOX so it cannot be related to Tbx5 expression levels. All together these data show that Tbx5 overexpression does not direct cardiac differentiation to the SAN fate.

### Blocking TAK1 phosphorylation activates SAN markers in cardiac differentiation

We previously showed that TAK1 overexpression directs cardiac differentiation toward the SAN fate (Brown et al. 2017). Here we show that Tbx5 overexpression increases TAK1 expression but does not increase SAN differentiation. This may be due to a dose effect however since we have also shown that EBs constitutively overexpressing TAK1 ultimately show decreased phosphorylation of known downstream targets (Hunter et al. 2019) it is possible that SAN differentiation is related to this downregulation. To test this, EBs were differentiated in the presence of OXO. OXO-treated EBs beat faster than untreated EBs and had a similar range of beat rates both as our original PGK::TAK1 EBs and the TAK1 inducible EBs (Fig.6A). OXO treatment also impacted the morphology of cardiac cells at day 21, (Compare Fig.6B to Fig.6C) with cells adopting a spindle-like morphology which is characteristic of SAN cells in vivo (Wu et al. 2001). To test whether individual cardiomyocytes activated the SAN transcriptional program, immunocytochemistry was performed to assess Islet1, and Shox2 expression. OXO treatment dramatically increased the percentage of cardiac cells expressing these SAN transcription factors as well as the percentage of cardiac cells expressing the SAN channel protein HCN4 (Fig.6C).

**Figure 6:**
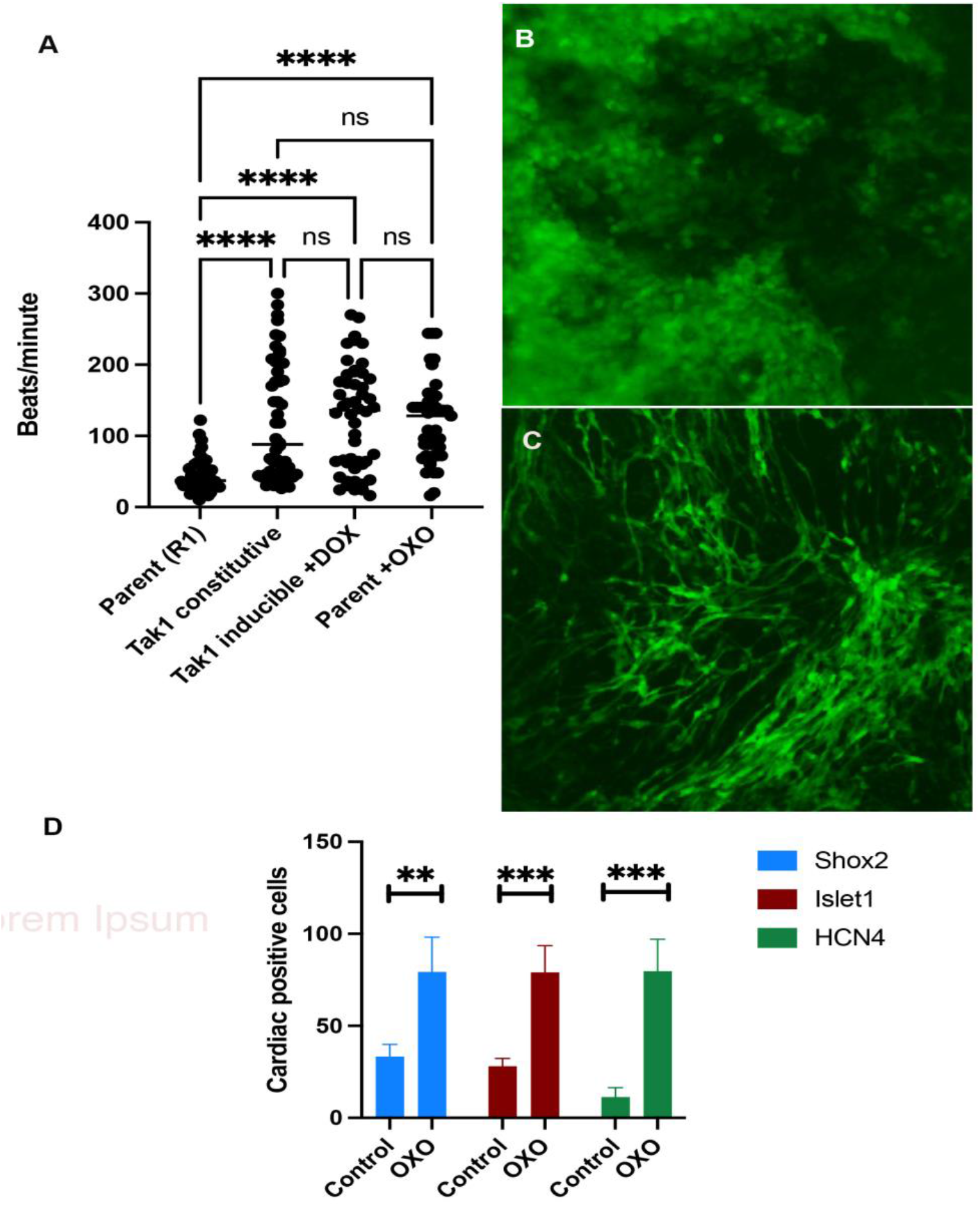
Blocking TAK1 phosphorylation increases expression of SAN transcription factors in cardiomyocytes. A) Beat Rate data day 21 cardiomyocytes from R1, TAK1 constitutive, TAK1-inducible with DOX, R1 with OXO, EBs. Statistical significance was determined by ANOVA. B) Live cell imaging of day 21 cardiomyocytes from R1 and (C) R1 with OXO treatment, imaging fluorescence from the αMHC:GFP cardiac promoter reporter. D)Quantification of Shox2, Islet1, and HCN4 expression in cardiac cells as assessed by ICC, with and without OXO treatment. Data represents means ± standard error of at least 3 independent experiments. Statistical significance was determined by an unpaired two-tailed t-test. **p<0.01, ***p<0.001, ****p<0.0001.

## Discussion

We produced mouse embryonic stem cell lines that conditionally overexpress Tbx5, and other factors involved in SAN differentiation. One of these cell lines, B1 increased the rate of beating in cardiac cells across a range of doses and showed increased expression of Tbx5 protein in response to DOX.

Tbx5 is initially expressed throughout the cardiac crescent but becomes restricted to the sinoatrial region during linear heart tube formation. Both homozygous and heterozygous loss of Tbx5 causes conduction system defects and loss of SAN structures (Bruneau et al. 2001, Moskowitz Ivan PG et al. 2004) demonstrating that it plays an important role in the development of the SAN. By contrast, when Tbx5 is ectopically mis-expressed in ventricular tissues, as was seen in the Mef2C-/- mouse (Vong et al. 2006) ventricular cells did not convert to an SAN-like fate but rather adopt an atrial fate. Together, these data suggest that overexpression may not be sufficient for cells to adopt the SAN fate. Our data appear to agree with this general finding as EBs overexpressing Tbx5 did not adopt indicators of the SAN fate.

Interestingly, our data demonstrate that Tbx5 overexpression resulted in the upregulation of the Map kinase, TAK1. We previously showed that TAK1 is expressed strongly in cells that will ultimately give rise to SAN in vivo and that EBs overexpressing it direct most cardiac cells to the SAN fate. However, we also showed that EBs with forced overexpression of TAK1 significantly downregulate the transcription of endogenous TAK1 (Brown et al. 2017) and the phosphorylation of known downstream targets, p38 and Jnk (Hunter et al. 2019). These data suggest that TAK1 expression levels are generally under strict transcriptional control. Our findings are consistent with in vitro studies demonstrating that TAK1, can inhibit the phosphorylation of its downstream targets in a context-dependent fashion (Ajibade et al. 2012, Cheung et al. 2003). To understand this thoroughly will require studies to determine which of TAK1’s known targets are impacted, and whether there are additional as yet unidentified (Ishitani et al. 2003, Ishitani et al. 1999, Moriguchi et al. 1997, Ninomiya-Tsuji et al. 1999, Shirakabe et al. 1997)TAK1 phosphorylation targets. For example, SAN fates can be activated in EBs treated with suramin which inhibits the phosphorylation of Jnk (Wiese et al. 2011). More recently, it has been demonstrated that post-translation modifications of TAK1 and Tab1 (Hirata et al. 2017) can impact the phosphorylation of downstream targets. The impact on TAK1 on SAN has not been studied in vivo because both the original gene trap of TAK1 (Jadrich et al. 2006) and studies knocking it out in the entire heart using a floxed allele (Xie et al. 2006) are mid-gestation lethal.

Activation of SAN-like cells through the activity of BMP signaling (Protze et al. 2017) may seem inconsistent with these findings however, it is known that the TAK1-p38 MAPK or JNK pathways can be regulated in positive or negative manners by inhibitory SMADs. For example, SMAD7 activates the TAK1-p38 MAPK pathway in human prostate cancer cells (Edlund et al. 2003)) whereas Smad6 inhibits TGF-β1-induced activation of this pathway (Jung S. M. et al. 2013). In addition, it is known that canonical BMP/SMAD signaling is temporally regulated during cardiac differentiation with activation required for the early cardiac differentiation but repression of those same pathways required for later specification of cardiac cell types (Guzzo et al. 2007).

## Acknowledgements

The authors would like to thank Andrew Hunter PhD for feedback on these studies and comments on the initial text. We would also like to thank Jacob Kendrick and Richard Visconti PhD for discussion and assistance with flow cytometry studies.

